# Cytotrophoblast cells are selectively permissive and favor Zika virus, but no other related flavivirus, invasion to the placental stroma

**DOI:** 10.1101/2023.01.20.524913

**Authors:** Mercedes Viettri, Gerson Caraballo, Ma. Elena Sanchez, Aurora Espejel-Nuñez, Abigail Betanzos, Vianney Ortiz-Navarrete, Guadalupe Estrada-Gutierrez, Porfirio Nava, Juan E. Ludert

## Abstract

**BACKGROUND:** Zika virus (ZIKV) is highly teratogenic, in contrast with dengue virus (DENV) or the yellow fever virus vaccine (YFV-17D). The mechanisms employed by ZIKV to cross the placenta need to be elucidated.

**METHODS:** Parallel infections with ZIKV, DENV and YFV-17D were compared in terms of efficiency, activation of mTOR pathways and cytokine secretion profile in human cytotrophoblastic HTR8 cells (CTB), and monocytic U937 cells, differentiated to M2 macrophages (M2-MØ).

**RESULTS:** In CTB, ZIKV replication was significantly more efficient than DENV or YFV-17D. In M2-MØ, ZIKV replication continued to be more efficient, although differences between strains were reduced. Significantly greater activation of Phospho-S6r and Phospho-AKT/Ser473 fractions in CTB infected with ZIKV than with DENV or YFV-17D, was observed. CTB treated with the mTOR inhibitors rapamycin or AZD8055, showed a 20-fold-reduction in ZIKV yield, versus 5 and 3.5-fold for DENV and YFV-17D, respectively. Finally, we detected that ZIKV infection, but not DENV or YFV-17D, efficiently inhibited the interferon response of CTB cells.

**CONCLUSIONS:** These results suggest that CTB cells are permissive and act favoring ZIKV entry into the placental stroma, over DENV and YFV-17D and that the mTOR complex is a switch that enhances the replication of ZIKV in CTB cells.

## 1. INTRODUCTION

The Zika virus (ZIKV) is a *Flavivirius*, mainly transmitted by mosquitoes^1,2^. Although, first isolated in 1947 in the Zika forests (Uganda), the first human outbreak was reported in 2007 on the Yap Island. In May 2015, the first cases in Brazil were reported and by 2016, ZIKV infections were reported by most countries in the Americas^1,3^. The ZIKV is classified into two genotypes, African and Asian; with the Asian lineage responsible for all the outbreaks in the Americas so far^4^.

ZIKV crosses the blood-placental barrier causing miscarriages and the congenital Zika syndrome (CZS), characterized by neurological disorders (microcephaly, anencephaly, and hydrocephalus), eye anomalies and congenital contractures^5,6^. Therefore, ZIKV is classified as a TORCH pathogen (*T. gondii*., Rubella, Cytomegalovirus, HIV, and other microorganisms capable of crossing the blood-placental barrier)^5^. In contrast, other mosquito-borne, phylogenetically, and epidemiologically related flavivirus, such as dengue virus (DENV) or the attenuated vaccine strain of yellow fever (YFV-17D) are not teratogenic^7,8^.

The main barrier of defense to avoid the vertical transmission of pathogens at the maternal-fetal interface is the chorionic villi, floating in close contact with the maternal blood in the intervillous space^9,10^. Chorionic villi are covered by two cell layers; the syncytiotrophoblast (STB) outer layer, formed from syncytialized trophoblasts in their terminal phase of differentiation, and the cytotrophoblast (CTB), an internal layer made up of trophoblasts that are still in the differentiation phase. These cell layers protect the placental stroma, where fibroblasts, Hofbauer cells (HBC), and fetal capillaries are found^9,10^. The STB is refractory to ZIKV infection due to secretion of interferon λ^11^. The rest of the microvilli cells, especially HBC, are susceptible to ZIKV infection^12,13^.

The mechanisms used by ZIKV to cross the blood-placental barrier are not well understood. One model proposes that during the first trimester of pregnancy, invasive trophoblasts encounter ZIKV-infected maternal cells of the basal decidua. Other routes proposed are that infected trophoblasts spread the virus from the decidua parietal across the amniochorionic membranes and release the virions directly into the amniotic fluid^14^, or the entry of ZIKV facilitated by cross-reactive DENV maternal antibodies through transcytosis^15^. Additionally, paracellular routes of invasion have also been suggested ^16^.

Regardless of the route used by ZIKV to enter the placental stroma, CTB cells should be involved in establishing the infection in the microvilli core; yet their role is unknown. In this work, ZIKV of African and Asian genotypes were studied in terms of infection efficiency, mTOR pathway activation, and cytokine response, using CTB and monocytes U937-DC-SIGN differentiated to M2-MØ, as trophoblastic and HBC placental cell models. DENV and the YFV-17D vaccine strains were compared side-by-side, given their similarities with ZIKV, but different teratogenic potential. The results suggest that the CTB plays a pivotal role in favoring the reach of the ZIKV to the placental stroma while deterring DENV and YFV-17D infections. In contrast, macrophages showed to be highly susceptible to all viruses and may act to amplify the ZIKV infection once inside the stroma. These results shed light on the mechanism of ZIKV vertical transmission.

## 2. MATERIALS AND METHODS

### 2.1 Cell lines

HTR8-SVneo cells, derived from human, first trimester of gestation placental trophoblast (ATCC®: CRL-3721), and U937-DC-SIGN cells, from human prohistiocytic lymphoma (ATCC®: CRL-3253), were grown in RPMI-1640 medium (Gibco®: 11875-093); Vero-E6 cells (ATCC®: CRL-1586) and mosquito C6/36 cells, derived from *Ae. albopictus* (ATCC®: CRL-1660), were grown in Eagle’s minimum essential medium (EMEM). All media was supplemented with 10% FBS and 100 U/ml penicillin-streptomycin. Mammalian cells were grown at 37°C and mosquito cells at 28°C, with 5% CO2.

### 2.2 Virus strains

The Zika virus strain Uganda (ZIKV-MR77), African genotype, was donated by Dr. Susana López, (IBT-UNAM, Cuernavaca). The Zika virus Mexican strain (ZIKV-MEX), Asian genotype, and the dengue virus serotype 2 strain New Guinea were provided by Dr. Mauricio Vázquez (InDRE, Mexico City). The YFV-17D vaccine strain was donated by Dr. Juan Salas-Benito (IPN, Mexico City). Viral strains were all propagated in C6/36 cells and titrated by focus assay in Vero-E6 cells.

### 2.3 Monocytes differentiation to M2 macrophages (M2-MØ)

U937-DC-SIGN cells were treated for 12 days with 160 nM Phorbol12-myristate-13-acetate (PMA) (Sigma-Aldrich®: P1585) and 30 ng/mL macrophage-colony stimulating factor (M-CSF) (Sigma-Aldrich®: M6518), replacing the medium every third day^17,18^. The percentage of differentiated monocytes was determined by flow cytometry and fluorescence microcopy.

### 2.4 One-step replication curves

Confluent monolayers of HTR8 cells and differentiated monocytes were seeded in 96-well plates and infected in triplicates, with ZIKV-MR77, ZIKV-MEX, DENV and YFV-17D strain using an MOI=3. Virus binding was carried out for 40 min at 4°C, to synchronize the infections. Afterward, cells were washed 3 times with cold PBS, maintenance media added, switched to 37°C and infections allowed to proceed for 0, 6, 18, 24, 48, 72 and 96 hours. At these times, supernatants were collected, and stored at -80°C until titration^19^. Briefly, Vero cells seeded in 96 well plates were infected with 10-fold serial dilutions of the supernatants. At 48hpi, cells were fixed with cold methanol, washed with PBS, and stained using anti-E Mab (4G2) as a primary antibody, and an anti-mouse IgG peroxidase conjugate (Jackson Immuno-Research®: 15-035-003) as secondary antibody. Infected cells were visualized using DAB (Vector-Laboratories®: SK-4100). Virus titers were expressed as FFU/mL.

In addition, the HTR8 and M2-MØ monolayers from the one-step replication curves were used to determine the percentage of infected cells at 24, 48, 72 and 96 hpi, by immunofluorescence. Briefly, monolayers were fixed in paraformaldehyde 4% for 10 min and permeabilized with 0.1% Triton X-100 for 10 min at room temperature. Cells were stained using a cross-reactive anti-NS3 Mab (1ED8) as primary antibody, and an anti-mouse Alexa-488 donkey pre-adsorbed (Abcam®: ab150064) as secondary antibody. Nuclei were counter stained with DAPI, and cells analyzed under a Nikon inverted microscope (Eclipse *Ti*-U). The virus yield/cell was calculated obtaining the ratio between viral progeny and the percentage of infected cells, assuming 10×10^5^ cells/well, after confluency.

### 2.5 mTOR activation in infected HTR8 cells

Activation of the mTORC1 and mTORC2 pathways was evaluated by western blot, analyzing the status of phospho-S6r for mTORC1 and phosphor-AKT-Ser473 for mTORC2. Infected, drug treated or control cells, were lysed in RIPA lysis buffer (5 M NaCl, 0.5 M EDTA pH 8, 10% sodium deoxycholate, 10% SDS, 10% triton X-100) with protease inhibitor cocktail (Sigma-Aldrich®: P8340). Protein concentration in the extracts was quantified using Bradford protein assay (BioRad®: 500-0006). Depending on the experiment, 15 or 50 μg of protein, were processed for 10% SDS-PAGE electrophoresis. Proteins were transferred to nitrocellulose membranes (BioRad®: 0.45 μm) following standard protocols. The following primary antibodies were used: a rabbit Mab S6 ribosomal protein (Cell Signaling®: 2217) and a rabbit Mab Phospho-S6 ribosomal protein (Ser235/236) (Cell Signaling®: 4858) for the mTORC1 pathway; a rabbit monoclonal antibodies AKT1 (Cell Signaling®: 2938) and Phospho-AKT (Ser473) (Cell Signaling®: 4060), proteins for the mTORC2 pathway. An anti-rabbit HRP (GeneTex®: GTX-26721) was used as a secondary antibody. All antibodies were diluted in 5% skim milk powder with PBS and 0.1% Tween-20. The HRP signal was detected using the SuperSignal™ West Femto kit (Thermo Fisher Scientific: 34096). Phosphorylated fractions were normalized with the same non-phosphorylated protein fraction. Digital images were captured with the ImageQuant LAS 4000 system (GE Healthcare) and analyzed with ImageJ software.

### 2.6 Inhibition of mTOR pathways in infected HTR8 cells

Rapamycin (Tocris-Bioscience®: 53123-88-9), was used to inhibit the mTORC1 pathway, and the ATP-competitive AZD8055 (AstraZeneca®: 2525), for the inhibition of mTORC1 and mTORC2 pathways. Cells were treated with 100 nM of rapamaycin or 10000 nM of AZD8055, diluted in dimethyl sulfoxide (Sigma-Aldrich®: D-2650), for 24 h before infection. These concentrations proved to be non-toxic to HTR8 cells (Supplemental figure 3). Afterward, cells were infected at a MOI=3 with each viral strain; supernatants were collected (24 hpi) for virus yield determination by focus forming assay. In addition, viral protein expression was detected using the anti-NS3 mouse Mab 1ED8, and β-tubulin (GeneTex®: GTX-101279) as load control. An anti-mouse HRP (Jackson Immuno-Research®: 15-035-003) was used as secondary antibody.

### 2.7 Cytokine and chemokine responses in infected cells

HTR8 cells and M2-MØ, were either mock-infected, or infected at a MOI=3 with ZIKV-MR77, ZIKV-MEX, DENV-2 and YFV-17D. Cell supernatants were collected at 6, 24, 48 and 72 hpi, and analyzed in triplicates for the presence of the following cytokines and chemokines IL-1β, IL-6, IL-8/CXCL8, IL-10, IL-15, TNFα, MCP-1/CCL-2, MIP-1α/CCL-3, MCP3/CCL-7, IP-10/CXCL10, VEGF, IFN-2α and IFN-Y using a commercial multiplex system (Bio-Plex® Multiplex Immunoassay System, HCYTOMAG 13K MERCK). Samples were processed following the producer indications, and cytokine concentrations determined in a Luminex X-200. Results were plotted as line charts using GraphPad Prism software, version 6.01.

### 2.8. Statistical analysis

Values obtained from all experiments were expressed as means±standard errors (one-way ANOVA) of at least three independent experiments. Statistical analyzes and graphs were performed with GraphPad Prism software, version 6.01.

## 3. RESULTS

### 3.1 Infection efficiency of ZIKV, DENV and YFV-17D vaccine in CTB and macrophages

To investigate the mechanisms used by ZIKV to cross the placenta, the efficiency of infection and replication of two ZIKV strains, belonging to the African and Asian genotypes, was evaluated and compared with DENV and YFV-17D strains. One step replication curves were carried out in two cell lines, as placental models. HTR8 cells were used a CTB models and U937 monocytes differentiated to M2-MØ as HBC model. The M2-MØ phenotype was confirmed through expression analysis of the differentiation markers CD14, CD163 and CD209^20,21^ (Supplemental Figures 2); more than 95% of the culture expressed CD14 and CD163, indicating a macrophage M2 phenotype. CTB were susceptible to infection by all the strains tested; yet, the virus progeny for the ZIKV-MR77 and ZIKV-MEX strains reached significantly higher titers than for DENV and YFV-17D strains. By 48 hpi, 1×10^7^ and 1.5×10^6^ FFU/mL were obtained for the ZIKV-MR77 and MEX strains respectively, more than 1 log higher than the titers (1×10^5^ FFU/ml) reached for DENV and the YFV-17D vaccine strains at the same time (Figure 1A). On the other hand, M2-MØ cells were highly susceptible to infection by all strains. In macrophages ZIKV-MR77 titers reached 1×10^9^ by 48 hpi; yet the differences in yield among strains, were reduced to less than 1 log (Figure 1B).

**Figure 1.**
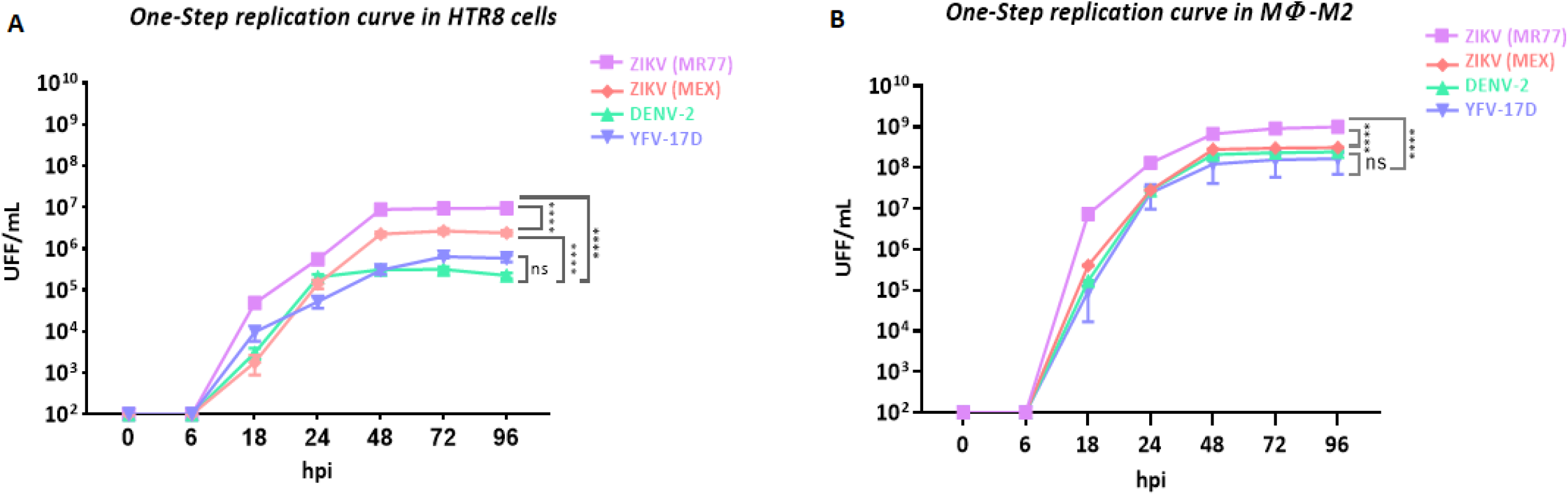
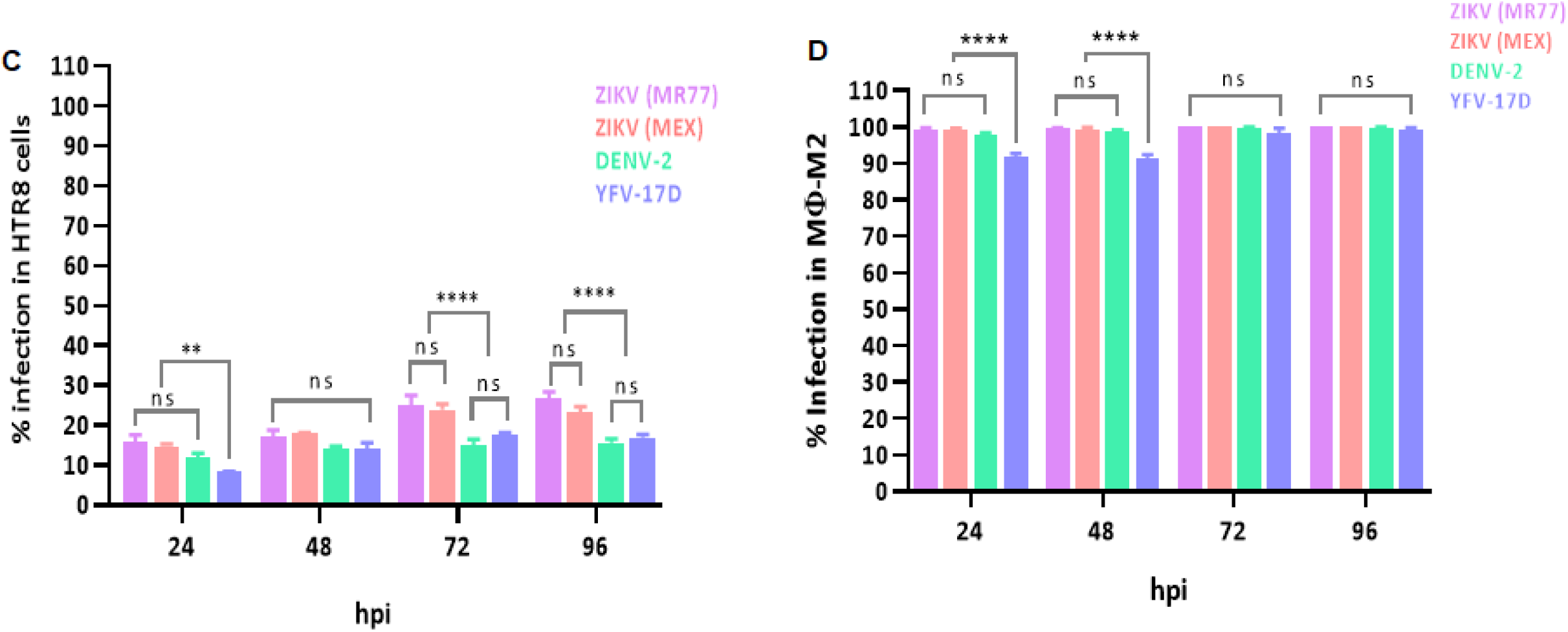
Replication efficiency of ZIKV-MR77, ZIKV-MEX, DENV-2, and YFV-17D vaccine strains in cytotrophoblastic cells and differentiated macrophages. A) One step replication curves in HTR8 and B) in M2-macrophages derived from U937 cells. Infections (MOI=3) were synchronized by incubation of the monolayers on ice. Supernatants were titrated in Vero-E6 cells using focus-forming unit assays. Curves showed mean titers±standard deviation of at least 3 independent experiments. C) Percentage of infected HTR8 cells. D) percentage of infected M2-macrophages. Infected cells were stained using a cross-reactive anti-NS3 Mab, and results expressed as percentage of positive cells in relation to the total number of cells stained with DAPI. n=3 for both cell lines. **** p ≤ 0.0001; ** p ≤ 0.0090; ns, not significant.

The higher ZIKV strains infection efficiency in HTR8 cells was corroborated when the percentage of infected cells (Figure 1C) or the amount of virus produced per cell was determined. ZIKV strains always showed a higher percentage of infected HTR8 cells than DENV or YFV-17D, with differences becoming statistically significant (25-24% versus 15-17%; p≤ 0.0001) at 72 hpi. In turn, macrophages were infected over 90% by all strains even at 24 hpi (Figure 1D). Virus yield per cell were calculated with the data obtained from the viral replication kinetics and the infection percentages (Table 1). Infected HTR8 cell produced ZIKV-MR77 viruses up to 20-fold times more than DENV or YFV-17D infected cells (275 versus 12 and 14; p≤0.0001). For the ZIKV-MEX strain, differences in virus yield per cell with DENV-2 and YFD-17D strains were less marked, reaching 5-fold at 48 hpi, but were still significant. Finally, macrophages were significantly more permissive for the ZIKV-MR77 than to the other strains, with 5-fold differences, for ZIKV-MEX, DENV-2 and YFV-17D, respectively, at 48 hpi.

The results indicate that CTB and M2-MØ are susceptible to infection by all the tested viruses. However, in HTR8 cells ZIKV shows significantly higher efficiency in terms of infection and replication, than DENV and YFV-17D. Although a trend was also observed in M2-MØ, differences between strains were less marked.

### 3.2. mTOR pathway activation in infected CTB

The mTOR signaling pathway is used by RNA viruses to promote replication and evade the antiviral response^22-28^. Thus, we evaluated the activation of mTORC1 and mTORC2 in CTB cells infected with ZIKV, DENV or YFV-17D. A faster and stronger overactivation of mTORC1 (Figures 2A, 2B) and mTORC2 (Figure 2C, 2D) in CTB cells infected with ZIKV strains, when compared with DENV and YFV-17D, was observed. In ZIKV-MR77 infected cells, the activation of mTORC1 and mTORC2 peaked at 6 and 12 hpi, respectively and doubled in ZIKV-MR77 infected cells when compared with DENV and YFV-17D. mTORC1 and mTORC2 activation in ZIKV-MEX infected cells was also significantly higher than the induced after DENV and YFV-17D infection but was reduced in comparison with ZIKV-MR77 (Figures 2C, 2D). Finally, the infection with DENV and YFV-17D activated the mTOR signaling at similar extent, about 0.5-fold over background levels (mock-infected cells). These results suggest that ZIKV is a potent inductor of the mTOR pathways in CTB cells; yet significant differences were observed between the African and Asian lineages.

**Figure 2.**
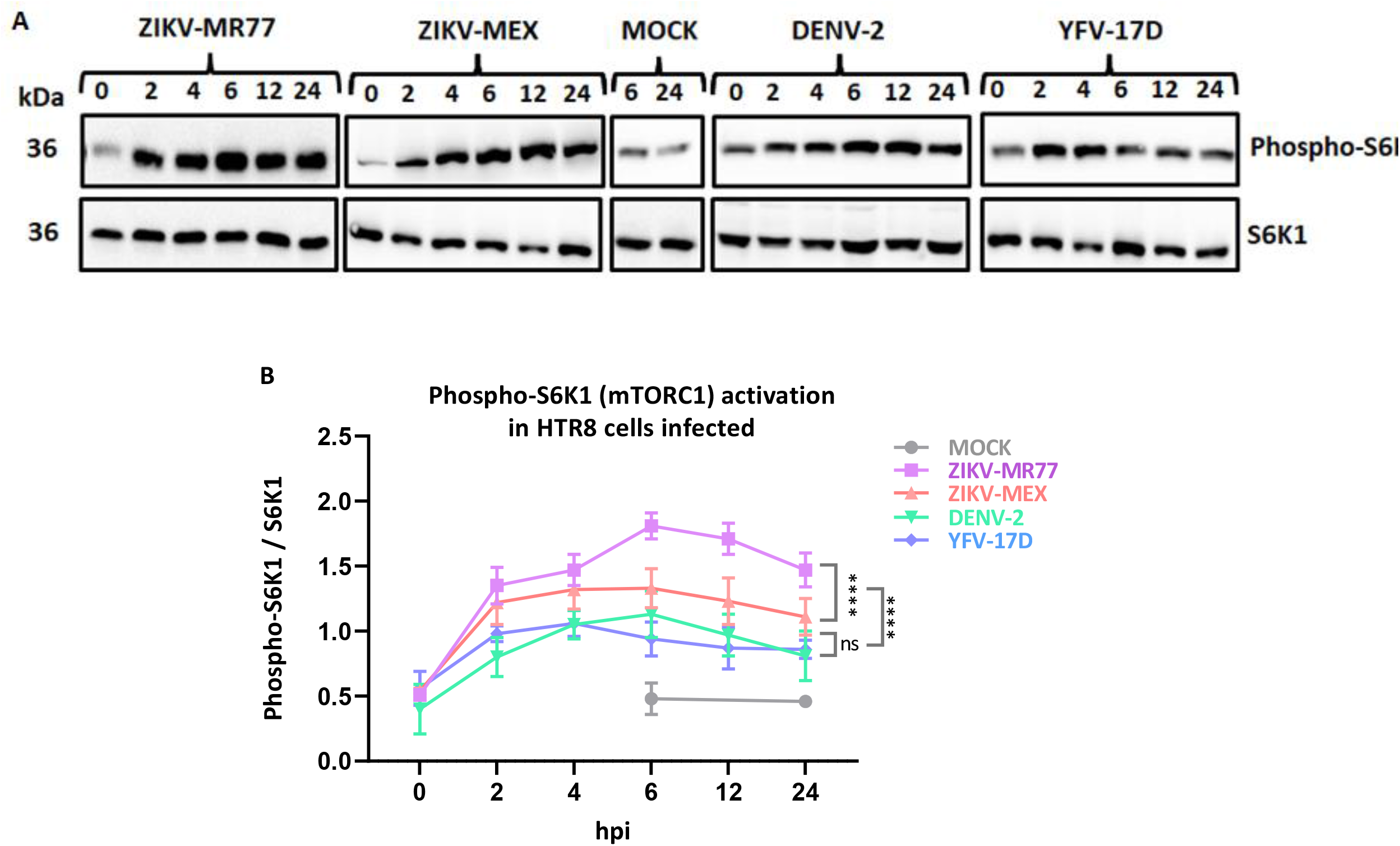
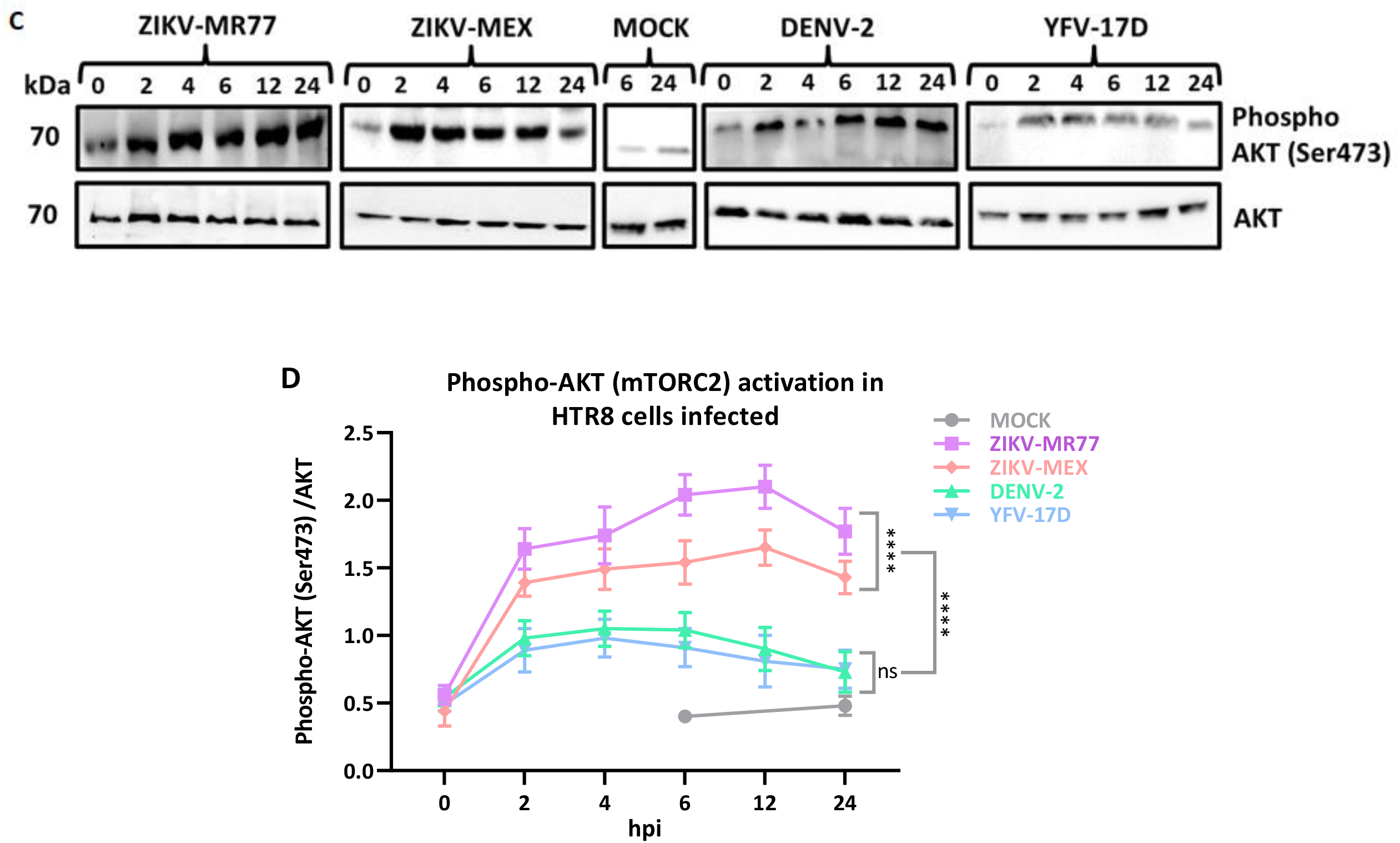
Activation of mTOR pathways in infected cytotrophoblastic cells. Confluent monolayers of HTR8 cells were infected with the indicated viruses (MOI=3), or mock infected. Cell lysates were collected at different hpi, analyzed by WB, and band intensities assessed by densitometry. Activation of the mTORC1 pathway was determined measuring the phospho-S6/S6 ratio (panels A and B), and the phospho-AKT/AKT ratio for the mTORC2 pathway (panels C and D). All experiments were carried out in triplicates, graph lines represent means ± standard errors. **** p≤0.0001.

### 3.3 Effect of rapamycin and AZD8055 treatment of infected CTB

Rapamycin and AZD8055, two allosteric inhibitors of the mTOR pathway ^29,30^, were used to determine the function of mTOR signaling in virus biogenesis. Both viral NS3 protein production and virus yield were evaluated at 24 hpi. CTB cells remain metabolically viable 48 h after treatment, even with the highest concentrations tested: 100 nM for rapamycin and 10000 nM for AZD8055 (Supplemental figure 3). Furthermore, treatments were specific given that rapamycin inhibited mTORC1 activation, without affecting mTORC2, while AZD8055, inhibited both pathways (Figures 3A, 3B, 3C). Both drugs significantly reduced NS3 protein levels (Figures 3A, 3D), but no differences were observed between rapamycin and AZD8055 treatments, indicating that the activation of mTORC1 partially mediates the viral replication. In addition, viral titers were significantly reduced in rapamycin and AZD8055 treated cells, with a greater reduction observed for ZIKV-MR77 (over 1 log reduction). No significant differences in virus titers were observed in cells treated with rapamycin or AZD5088, suggesting that it is the mTORC1 pathway which is mainly required for viral replication (Figure 3E). Importantly, DENV and YFV-17D replication was also mTORC1 dependent (Figure 3).

**Figure 3.**
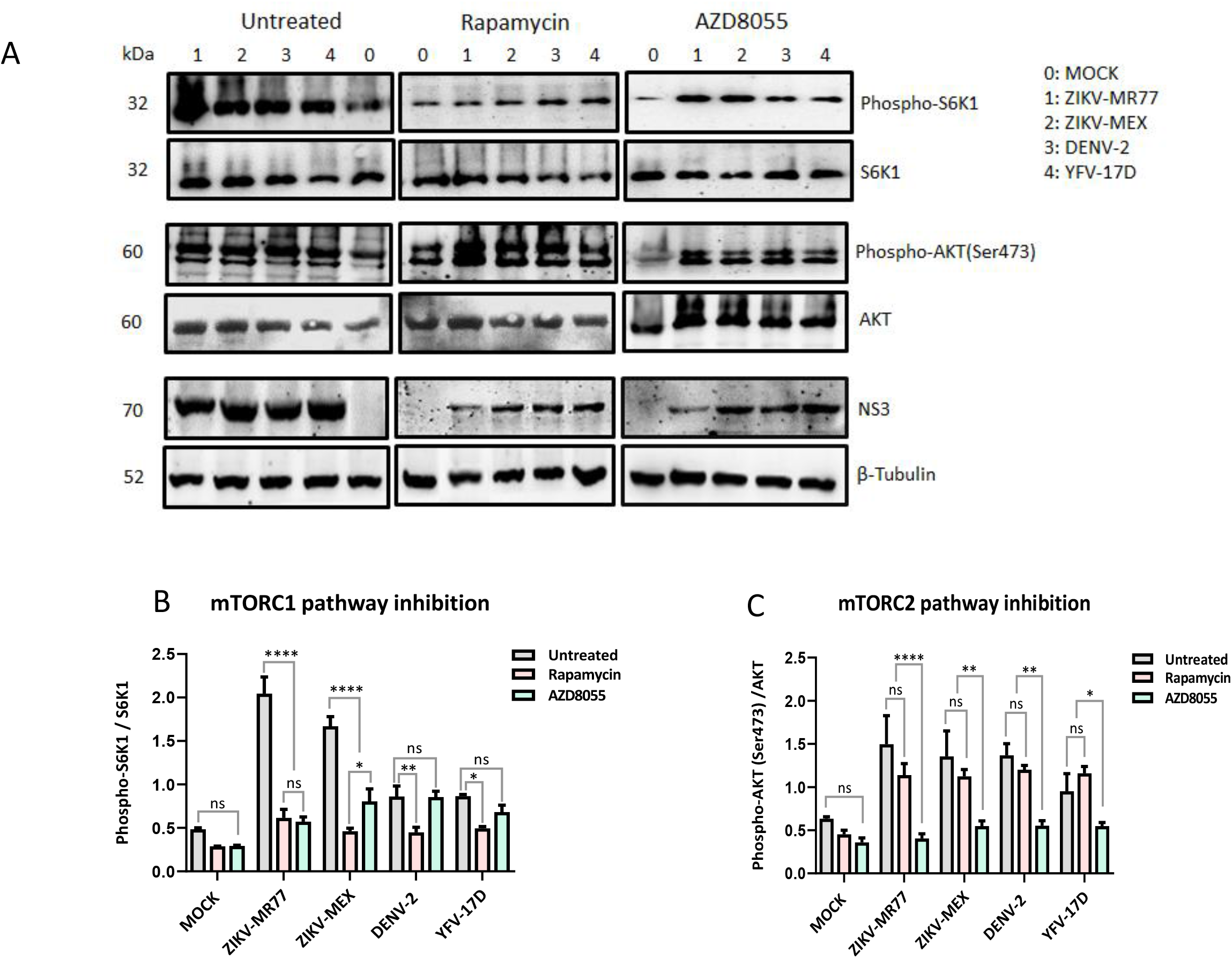
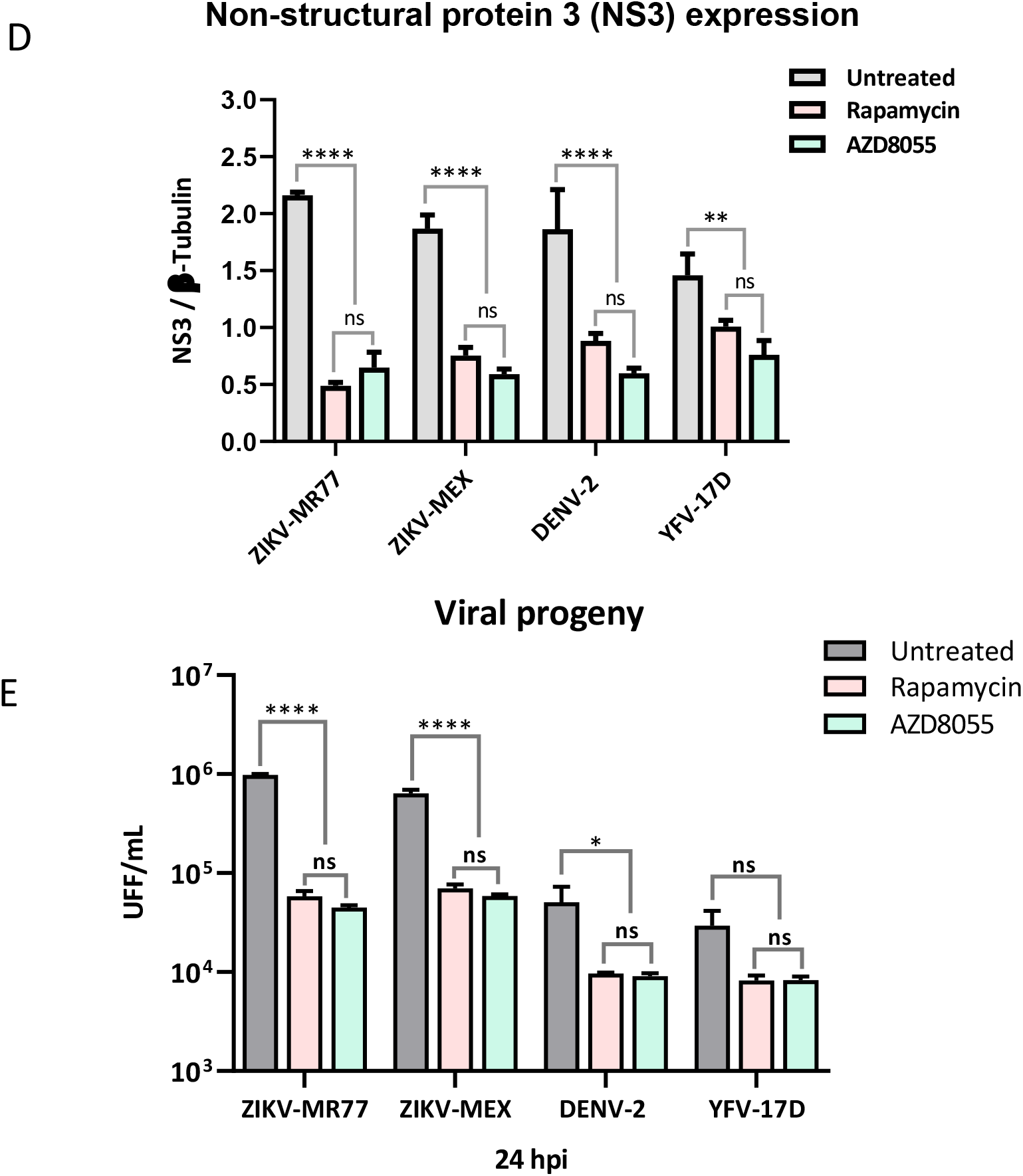
Effect of the inhibition of the mTOR pathways on virus replication. HTR8 cells were pretreated with rapamycin or AZD8055, for 24 h before viral infection. Twenty-four hours post infection, cells were lysed, analyzed by WB (panel A) and band intensities assessed by densitometry to assess the drug effect on mTORC1 (panel B), mTORC2 (panel C) inhibition, and NS3 expression levels (panel D). Supernatants were analyzed by focus assay for virus titers determinations (panel E). Bars show means ± standard deviations of at least 3 independent experiments. **** p≤0.0001; ** p≤0.0065; *p≤0.0229; ns, not significant.

### 3.4 Cytokine and chemokine responses in infected CTB and macrophages

The delicate balance between the anti- and inflammatory responses is crucial for the development of the semi-allogeneic fetus^9,10^. All tested cytokines were induced after infection by ZIKV-MR77, ZIKV-MEX, DENV and YFV-17D strain in both, CTB and M2-MØ. The induction was more evident at later time points (24-72 hpi) and more robust in M2-MØ (Figures 4 and 5; supplemental tables 2 and 3). The cytokine profile of CTB cells infected with ZIKV strains was characterized by high levels of IL-1β, IL-6, TNFα and low levels of IL-10, CCL-3, IP-10, VEGF, IFN-2α and IFN-γ (Figures 4A, 4B, 4C). Differences between ZIKV strains and DENV and YFV-17D were particularly evident for CCL-3, IP-10, IFN-2α and IFN-γ, suggesting that in CTB cells the infection with ZIKV negatively modulates the production of chemoattractant and recruiting chemokines, and IFNs.

**Figure 4.**
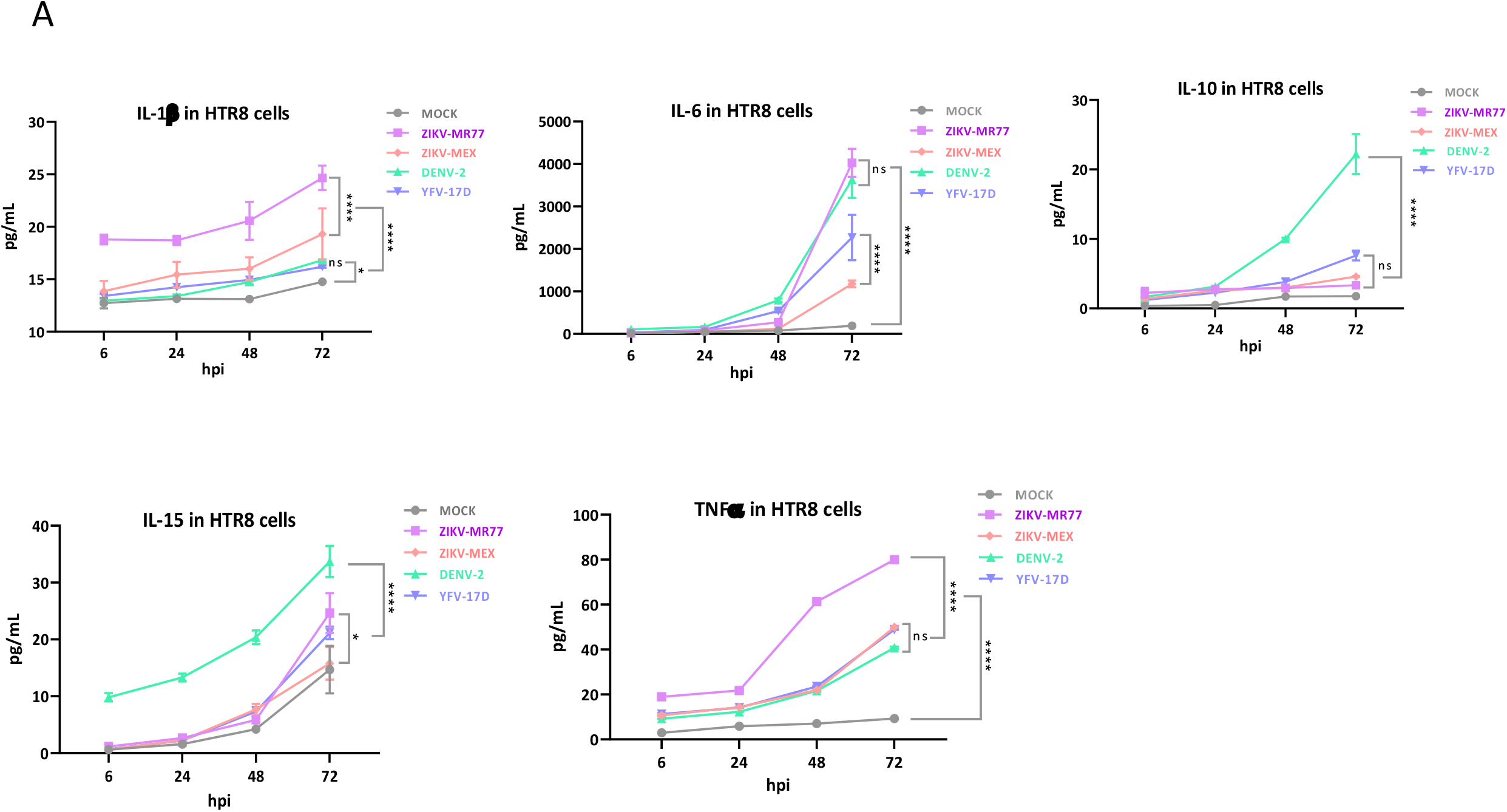
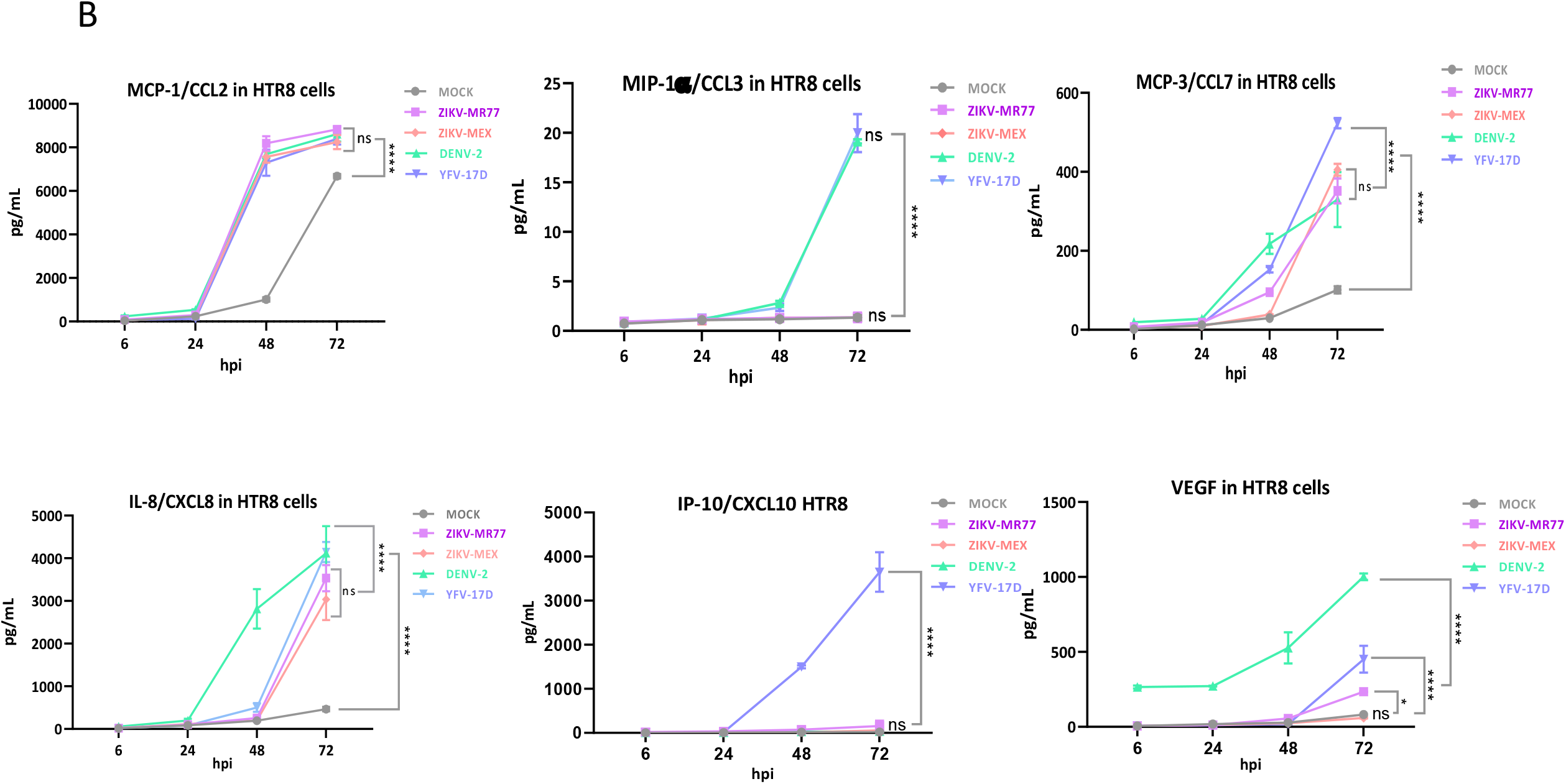
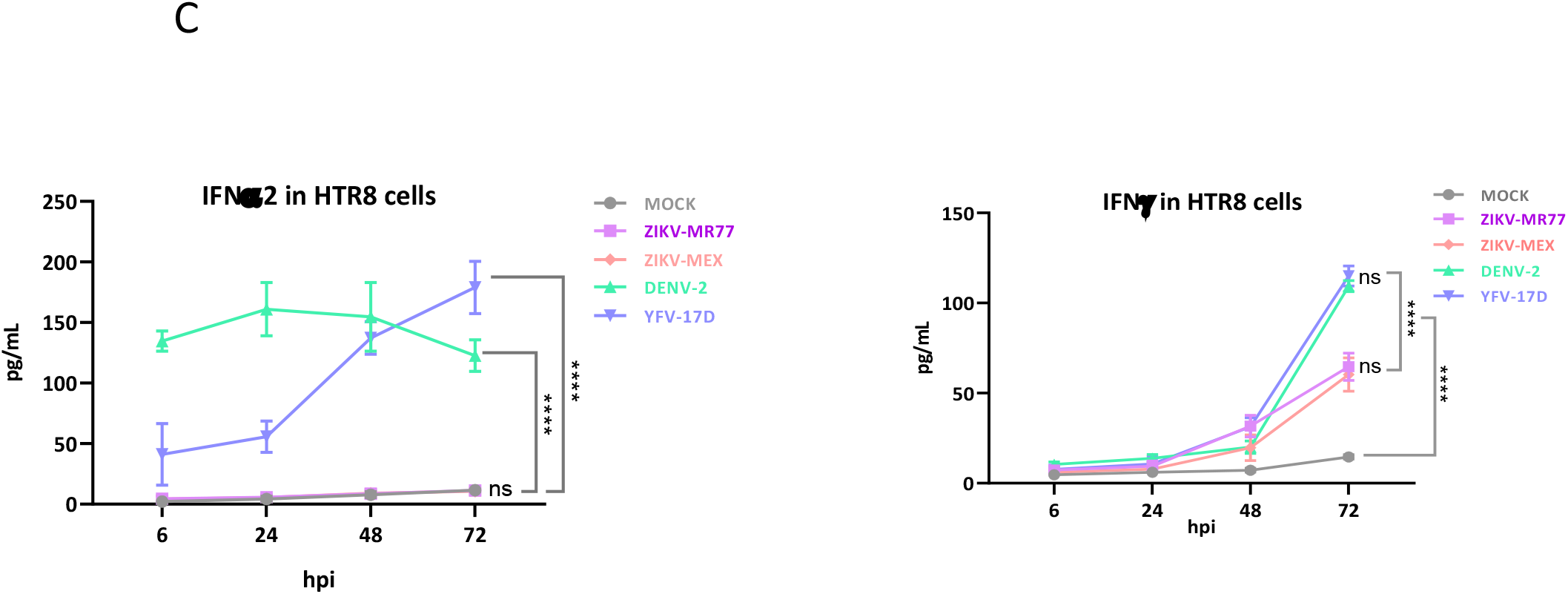
Cytokine profiles produced by infected CTRs cells. Confluent monolayers of HTR8 cells seeded in 24 well-plates were infected (MOI=3) with the different virus strains and supernatants collected at the indicated times. The concentration of secreted cytokines, chemokines and INF were determined by cytometric bead arrays. Graphs show mean concentrations (pg/ml) ± standard deviations of at least independent experiments. ****p, ≤ 0.0001; *p, ≤ 0.046; ns, not significant.

Interestingly, except for CCL-7, the differences in cytokine expression triggered by the ZIKV strains and DENV and YFV-17D strains were less evident in M2-MØ. Nonetheless, the ZIKV-MR77 generated a higher inflammatory and chemotactic response with superior values of IL-1β, IL-6, TNFα, CCL-2, CCL-3, CCL-7, CXCL8, IP-10 and VEGF, than the rest of the strains (Figures 5A, 5B). The poor IFN response induced by ZIKV strains was also observed in M2-MØ (Figure 5C).

**Figure 5.**
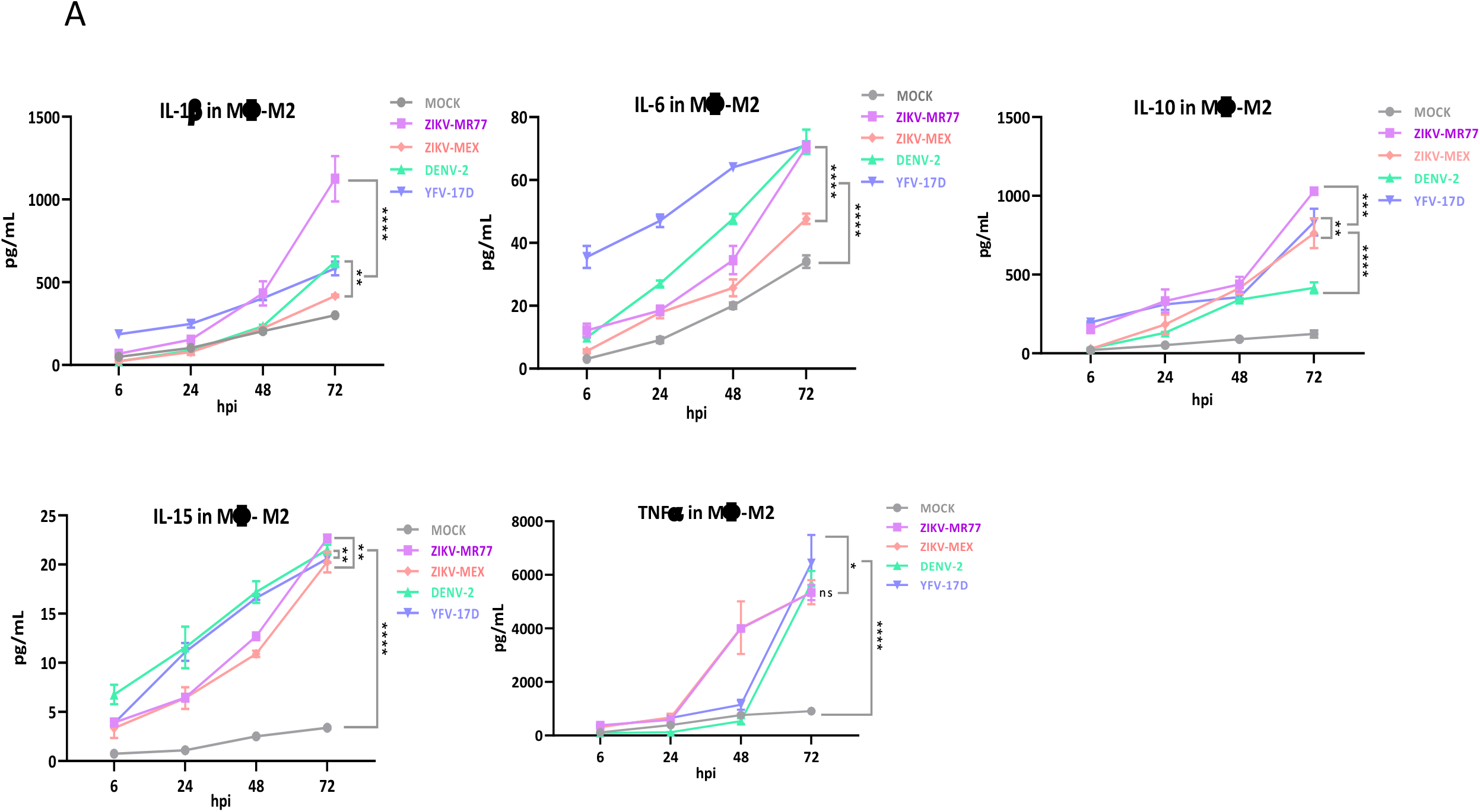
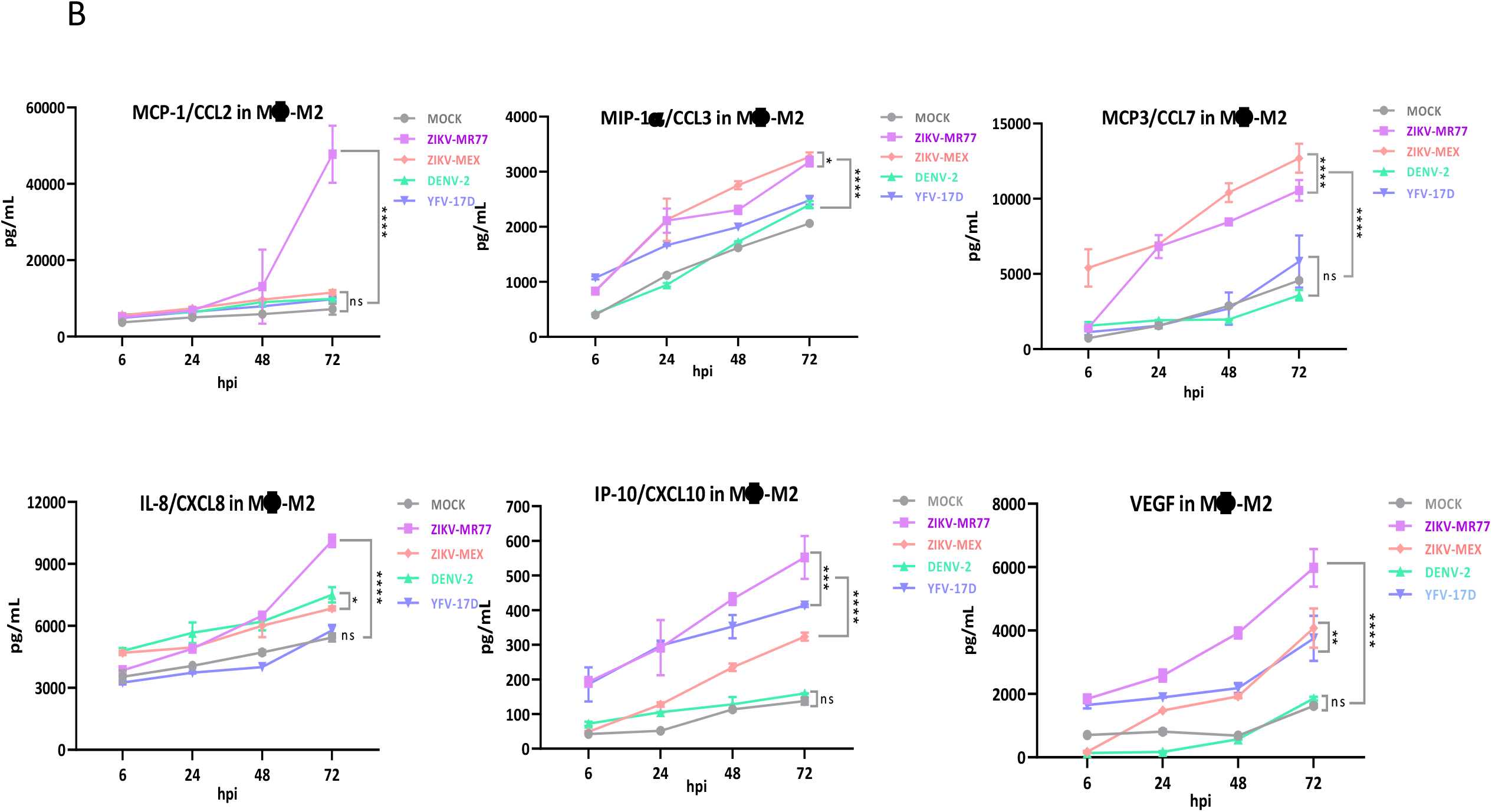
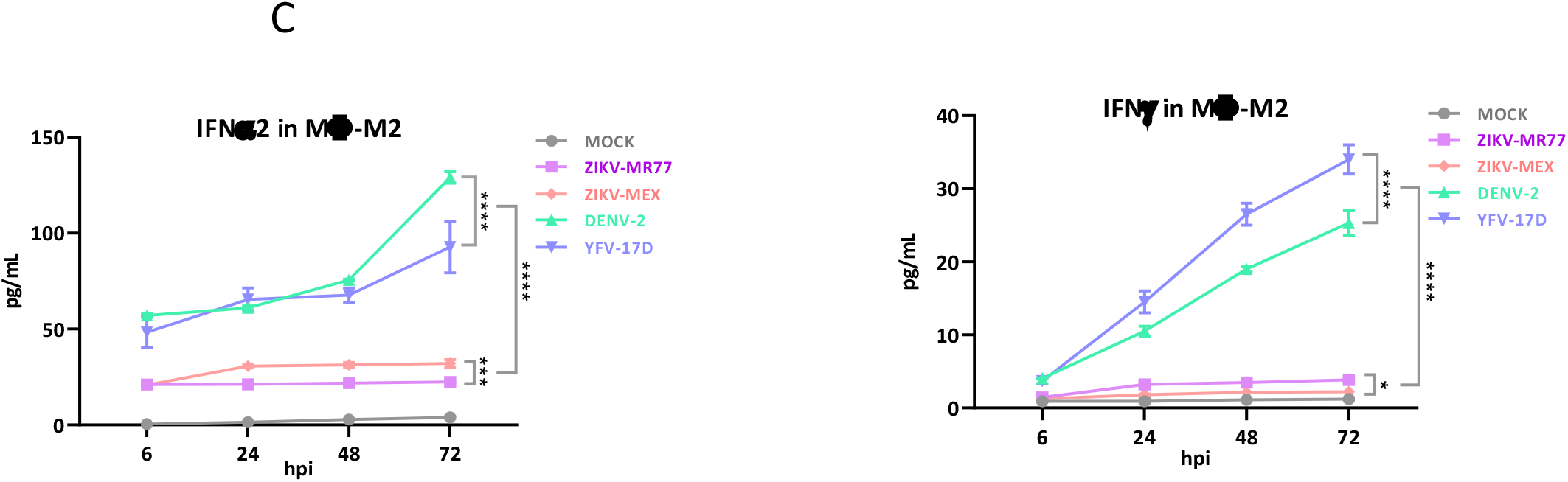
Cytokine profiles produced by infected differentiated M2 macrophages. Confluent monolayers of differentiated macrophages seeded in 24 well-plates were infected (MOI=3) with the different virus strains and supernatants collected at the indicated times. The concentration of secreted cytokines, chemokines and IFN were determined by cytometric bead arrays. Graphs show mean concentrations (pg/ml) ± standard deviations of at least independent experiments. ****p ≤ 0.0001; ***p ≤ 0.0008; **p ≤ 0.0056; *p, ≤ 0.0461; ns, not significant.

## 4. DISCUSSION

ZIKV is a TORCH pathogen with high teratogenic potential^5^; in contrast, other related flaviviruses, such as DENV and YFV, have never been associated with fetal damage^2,7,8^. How the ZIKV cross the blood placental barrier need to be deciphered, given the social and public health impact generated by the CZS. Parallel infections using African and Asian ZIKV strains, along with DENV and YFV-17D, highlighted the role of CTB cells in gating the pass of ZIKV, but not DENV or YFV, to the placental stroma. In agreement with previous results, infection assays indicated that HTR8 cells, despite the absence of DC-SIGN expression (Supplemental figure 2D), are susceptible to ZIKV, DENV and YFV ^31^. However, both ZIKV strains, particularly the African strain, showed significantly higher infection and replication capacity in HTR8 cells than DENV and YFV-17D. On the other hand, MØ-M2 were highly permissive to all four strains, and the marked differences observed in HTR8 cells, were reduced, most likely reflecting the high susceptibility of cells of the monocytic lineages, to all mosquito-borne flavivirus^32,33^. HBC, that have been implicated in the vertical transmission of ZIKV^12,13^. Our results suggests that once inside the placental stroma, ZIKV infection is amplified by HBC, favoring virus spread to the fetal circulation^13,14^.

Congenital malformations associated with ZIKV have been attributed only to the Asian lineage^3,4^. However, our findings agree with previous reports using human placental trophoblast, animal cell lines, mosquitoes and mice models, which all evidenced the higher capacity for transmission, replication, and virulence of the African over the Asian lineage^34,35^. Thus, African strains shall be seen as virulent, which could generate early miscarriages, preventing advanced fetal development^36^.

Cell permissiveness to a viral infection is the result of multiple factors. The mTOR machinery is a cell signaling pathway related to the regulation of various processes such as growth, proliferation, survival, protein synthesis, ribosome and lipid biogenesis, autophagy, and migration^37,38^. This pathway has been reported to be essential during replication and induction of antiviral immune responses by various RNA viruses including ZIKV, DENV, and YFV^22-28,39^. Our results suggest that in HTR8 cells, ZIKV, especially the African strains, activate mTOR signaling far more effectively than DENV and YFV-17D. Inhibition by chemical means of mTOR signaling, resulted in decreased NS3 viral protein expression and lower the viral titers in all infection conditions. The stronger inhibition in the replication of ZIKV after rapamycin and AZD8055 treatment demonstrated a higher reliance of the ZIKV strains in the activation of mTOR, and specifically in the activation of mTORC1.

Our results agree with and expand previous works reporting the activation and the dependency on the activation of the AKT/mTORC pathways of ZIKV infections, using liver, neuroblastoma, neural progenitor, and embryonic kidney human cell lines^23,25,27,28^. In all those relevant cell models, inhibition of the mTOR or PI3K/AKT pathways adversely affects ZIKV replication. mTORC1 activation in ZIKV infected cells have been associated with negative regulation of autophagy^23-25,27^ and enhanced NS5 polymerase activity to facilitate viral replication^28^. In addition, general processes such as viral protein synthesis^27,28,37-39^ and ER membrane remodeling^39^, are also presumably favored by mTOR modulation. In agreement, mTOR signaling inhibition in BHK-21 cells, reduced YFV infection^22^.

However, Liang *et al*., (2016)^40^ observed not activation, but inhibition of the AKT/mTOR pathway in ZIKV (African lineage) infected human fetal neural stem cells, leading to defective neurogenesis and aberrant autophagy, although the effect on virus replication was not directly evaluated. Conflicting results regarding mTOR activation have been also observed for cells infected with DENV. Inhibition of mTOR signaling in DENV infected HepG2^41^, BHK-21 and mouse neuroblasts^42^, HUVEC^26^ and in megakaryocytes cells^43^, accelerated the infection. These differences may obey to the fact that contrary to ZIKV infection, DENV infection seems to be favored by autophagy^26,41^.

Many works infer that the innate immune response contributes to the vertical transmission of the Zika virus and the development of SCZ^44-46^. The results in HTR8 cells indicate that ZIKV infection negatively modulates the chemotactic and interferon responses in CTB. Similar results using the same cell line were reported by Luo *et al*., (2018)^31^; higher levels of IL-6 and TNFα and lower levels of CCL3 and IFNs α/β secretion were observed for ZIKV infection than for DENV-4 and YFV-17D. Additionally, in another study conducted in human brain astrocytes, the high rate of viral replication of both African and Asian strains of ZIKV directly related with a limited response of proinflammatory and chemotactic cytokines (IL-6, IL-8, IL-12, CCL2, CCL5, IP-10)^46^.

The innate response was more robust in macrophages than in HTR8 cells, and the differences between ZIKV strains and DENV and YFV-17D, were less noticeable. Infection with both ZIKV lineages was characterized by a more proinflammatory and chemotactic profile, mediated by high levels of IL-1β, IL-6, TNFα, CCL-2, CCL-3, CCL-7, CXCL8, IP-10 and VEGF. Yet the IFNs α/y response was equally reduced in macrophages infected with ZIKV, compared to the DENV and YFV-17D strains. The ability of ZIKV to negatively modulate the interferon response has been reported^47-49^. Proposed mechanisms to explain the low IFN induced by ZIKV infections include proteasomal degradation of the STAT2 activator^48^ and modulation of the biogenesis of peroxisomes, the signaling platforms during the IFN response^49^. If these mechanisms operate in ZIKV infected HTR8 cells need to be investigated. The cytokine profiles observed in this work suggest that the establishment of ZIKV infection in the placental stroma is favored by the limited inflammatory, chemotactic response and the low levels of IFNs α/y induced in CTB, thus positively modulating their permissibility to ZIKV replication. Likewise, the low IFN response in macrophage cells may be decisive in the propagation of ZIKV to the fetal circulation.

In summary, these results suggest that CTB constitute a physical barrier that contain DENV and YFV infection, but is successfully surpassed by ZIKV. Once inside the placental stroma, ZIKV will target HBC cells, where infection is amplified, subsequently spreading to the fetal circulation. The capacity of the ZIKV to cross the CTB barrier appears to be the result of a sum of factors, including greater infection and replication efficiency, better capacity to activate the mTOR machinery and a limited inflammatory and chemotactic response with low IFN activity. (WORD TEXT COUNT: 3497).

## Acknowledgments

Authors want to thank Dr. Ferdinando Liprandi (IVIC, Caracas) for donating Mab 4G2, Dr. Eva Harris (UC Berkeley) for donating Mab 1ED8 and Dr. Patricia Talamás-Rohana for her critical reading of the manuscript.

## A conflict of interest statement

Authors declare no conflict of interest.

## A funding statement

This work was partially funded by ICGEB (Italy), grant CRP/MEX 17-02; and CONACYT (Mexico), grant Pronaii 302979 309 and A1-S-9005.

## Mention of any meeting(s) where the information has previously been presented

These results were partially presented in the IUMS Congress (2022), and in the XII Congress (2021) of the Mexican Society for Virology.

## Supplemental figure legends

**Supplemental figure 1. Monocyte differentiation determined by epifluorescence microscopy**. U937-DC-SIGN cells were grown in 24-well plates containing glass coverslips and treated with PMA and M-CSF at the concentrations described above during 3, 6 and 9 days; untreated U937-DC-SIGN were included as controls. At the indicated times, cells were washed once with PBS, fixed in paraformaldehyde 4% for 10 min, and permeabilized with 0.1% Triton X-100 for 10 min at room temperature. Cells were stained using a Mab CD-163 (Abcam®: ab182422), as primary antibody, and an anti-rabbit Alexa-594 donkey pre-adsorbed (Abcam®: ab150064) as secondary antibody. Coverslips were mounted in Fluoroshield™ with DAPI (Sigma-Aldrich®: F6057) and analyzed under an inverted Nikon microscope (model Eclipse Ti) (Scale bar 20 μm).

**Supplemental figure 2. Percentage of differentiated monocytes determined by cell flow cytometry**. Monocytic U937-DC-SIGN cells grown in suspension were treated at 0, 3, 6, 9 and 12 days with PMA and M-CSF; afterwards, cells were fixed in paraformaldehyde 4% for 5 min at room temperature and washed twice in phosphate buffer saline (PBS). Fixed cells were incubated with PBS and 5% FBS for 10 min in ice, to block antibody unspecific binding. Finally, cells were stained for three characteristic markers of macrophages M2: CD14, using an APC anti-CD14 mouse monoclonal (Abcam®: 60901); CD163, using a FITC anti-CD163 mouse monoclonal (BD Bioscience®: 563697), and CD209 (DC-SIGN), using a PE anti-DC-SIGN mouse monoclonal (Abcam®: 136333). All antibodies were diluted 1:100 in a final volume of 50 μL. For all conditions, cells were incubated with the antibody mixtures in ice for 1 hour, in the darkness and washed twice in PBS. Appropriate unstained controls were included for each antibody. Stained cells were analyzed in a flow cytometer BD-LSR II Fortessa, and at least 10,000 events were recorded. Graphics made with the software FlowJo v10.6.1. **A**. Analysis of SSC-A (cytoplasmic complexity) versus FSC-A (size). **B**. CD14 expression from in U937-DC-SIGN positive cells at 0, 3, 6, 9 and 12 days of differentiation. **C**. CD163 expression from in U937-DC-SIGN positive cells at 0, 3, 6, 9 and 12 days of differentiation.

**Supplemental figure 3. Evaluation of DC-SIGN expression in HTR8 cells**. HTR8 were fixed in paraformaldehyde 4% for 5 min at room temperature and washed twice in phosphate buffer saline (PBS). Fixed cells were incubated with PBS and 5% FBS for 10 min in ice, to block antibody unspecific binding. Finally, cells were stained for CD14, using an APC anti-CD14 mouse monoclonal (Abcam®: 60901); CD163, using a FITC anti-CD163 mouse monoclonal (BD Bioscience®: 563697), and CD209 (DC-SIGN), using a PE anti-DC-SIGN mouse monoclonal (Abcam®: 136333). All antibodies were diluted 1:100 in a final volume of 50 μL. For all conditions, cells were incubated with the antibody mixtures in ice for 1 hour, in the darkness and washed twice in PBS. Appropriate unstained controls were included for each antibody. Stained cells were analyzed in a flow cytometer BD-LSR II Fortessa, and at least 10,000 events were recorded. Graphics made with the software FlowJo v10.6.1.

**Supplemental figure 4**. Cell viability assay in HTR8 cells treated with rapamycin and AZD8055. Cells were seeded in 96-well plates, then treated with rapamycin and AZD8055 at the indicated concentrations, including non-treatment and DMSO, control conditions. Plates were incubated with the drugs for 48 hours. Cell viability was determined using Cell Titer 96 AQueous nonradioactive cell proliferation assay (MTS assay) (Promega®: G3580) used according to the manufacturer’s procedures. n=3.

## Notes

### Competing Interest Statement

The authors have declared no competing interest.

