## supplemental material for "Cytotrophoblast cells are selectively permissive and favor Zika virus, but no other related flavivirus, invasion to the placental stroma"

### Supplemental Figures

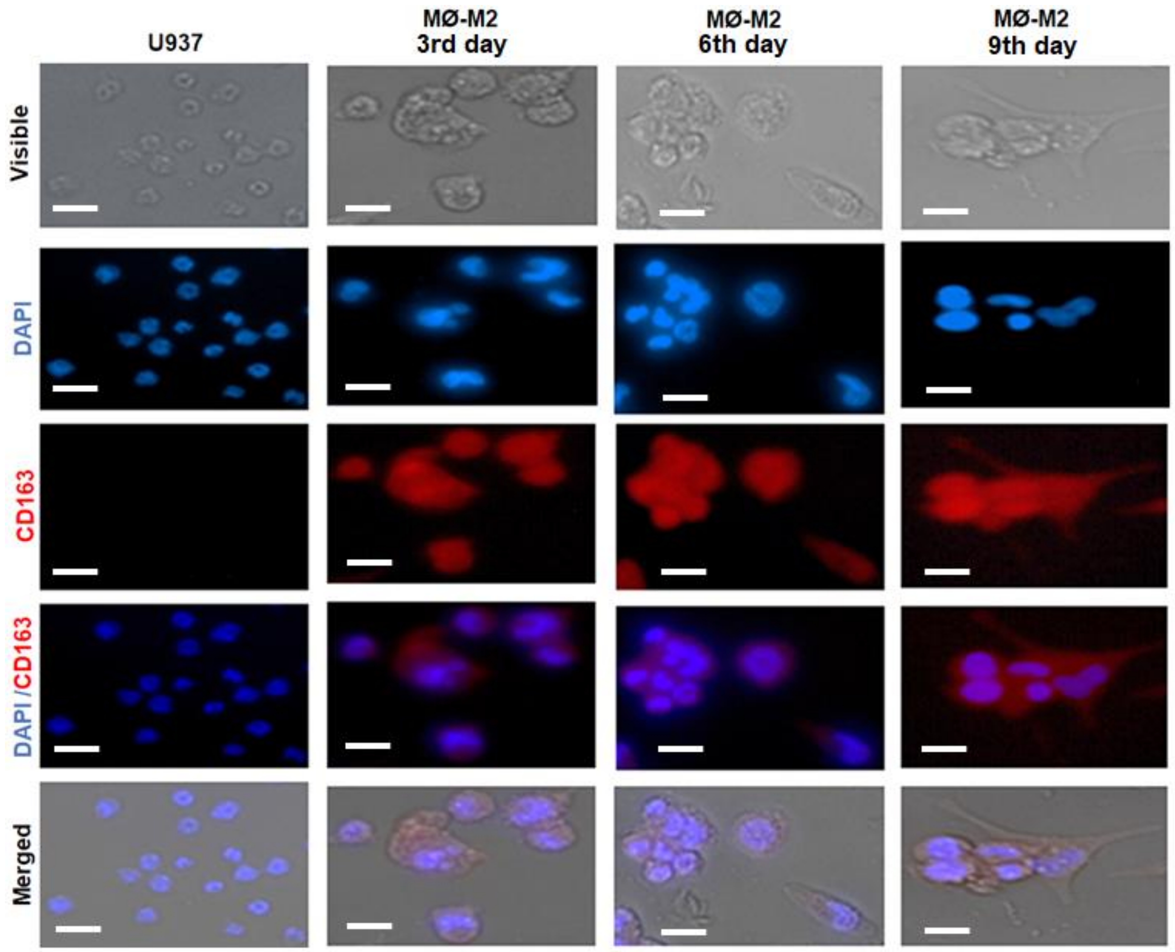


**Supplemental figure 1. Monocyte differentiation determined by epifluorescence microscopy.** U937-DC-SIGN cells were grown in 24-well plates containing glass coverslips and treated with PMA and M-CSF at the concentrations described above during 3, 6 and 9 days; untreated U937-DC-SIGN were included as controls. At the indicated times, cells were washed once with PBS, fixed in paraformaldehyde 4% for 10 min, and permeabilized with 0.1% Triton X-100 for 10 min at room temperature. Cells were stained using a Mab CD-163 (Abcam®: ab182422), as primary antibody, and an anti-rabbit Alexa-594 donkey pre-adsorbed (Abcam®: ab150064) as secondary antibody. Coverslips were mounted in Fluoroshield™ with DAPI (Sigma-Aldrich®: F6057) and analyzed under an inverted Nikon microscope (model Eclipse Ti) (Scale bar 20 µm).

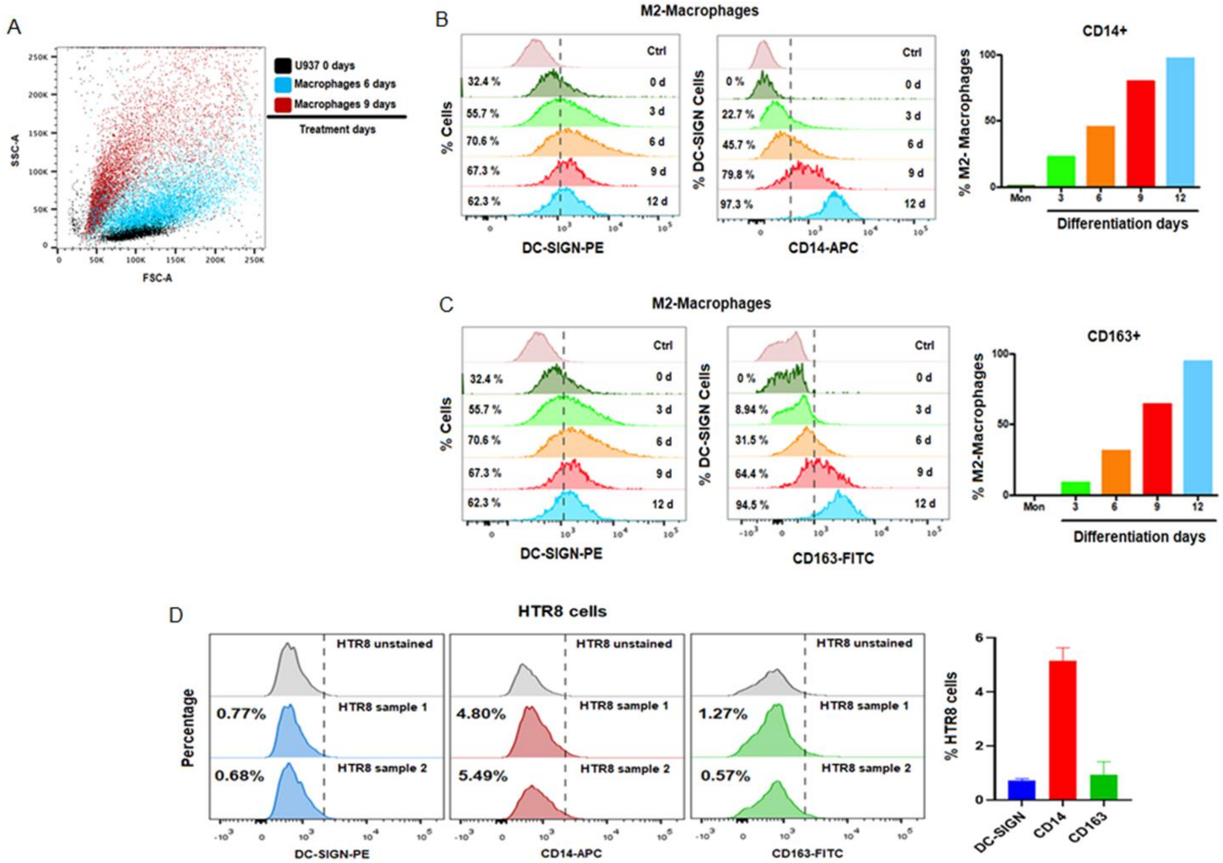

**Supplemental figure 2. Expression of DC-SIGN, CD14 and CD163 receptors in MØ-M2 and HTR8 cells.** Monocytes U937-DC-SIGN differentiated into M2-MØ and HTR8 cells were fixed in suspension with paraformaldehyde 4% for 5 min at room temperature and washed twice in phosphate buffer saline (PBS). Fixed cells were incubated with PBS and 5% FBS for 10 min in ice, to block antibody unspecific binding. Finally, cells were stained for three characteristic markers of macrophages M2: CD14, using an APC anti-CD14 mouse monoclonal (Abcam®: 60901); CD163, using a FITC anti-CD163 mouse monoclonal (BD Bioscience®: 563697), and CD209 (DC-SIGN), using a PE anti-DC-SIGN mouse monoclonal (Abcam®: 136333). All antibodies were diluted 1:100 in a final volume of 50 µL. For all conditions, cells were incubated with the antibody mixtures in ice for 1 hour, in the darkness and washed twice in PBS. Appropriate unstained controls were included for each antibody. Stained cells were analyzed in a flow cytometer BD-LSR II Fortessa, and at least 10,000 events were recorded. Graphics made with the software FlowJo v10.6.1. **A.** Analysis of SSC-A (cytoplasmic complexity) versus FSC-A (size). **B.** CD14 expression from in U937-DC-SIGN positive cells at 0, 3, 6, 9 and 12 days of differentiation. **C.** CD163 expression from in U937-DC-SIGN positive cells at 0, 3, 6, 9 and 12 days of differentiation. **D**. Expression of DC-SIGN, CD14 and CD163 in HTR8 cells.

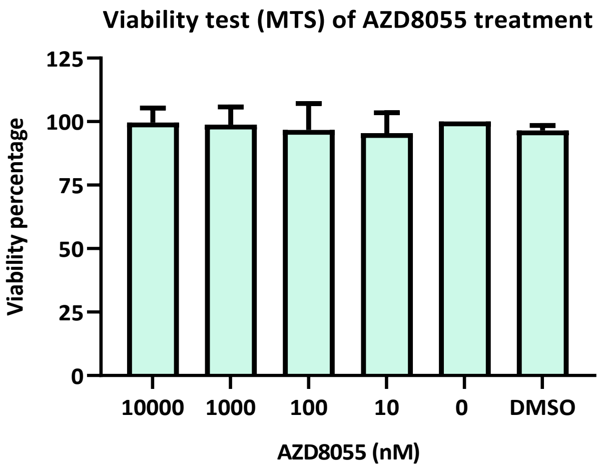


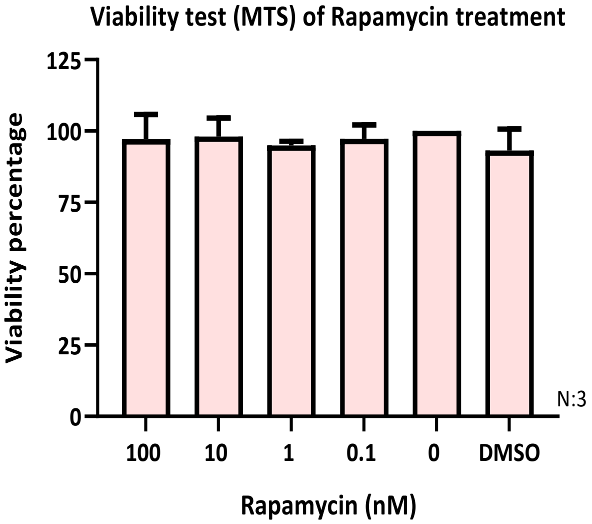


**Supplemental figure 3. Cell viability assay in HTR8 cells treated with rapamycin and AZD8055.** Cells were seeded in 96-well plates, then treated with rapamycin and AZD8055 at the indicated concentrations, including non-treatment and DMSO, control conditions. Plates were incubated with the drugs for 48 hours. Cell viability was determined using Cell Titer 96 AQueous nonradioactive cell proliferation assay (MTS assay) (Promega®: G3580) used according to the manufacturer’s procedures. n=3.

**Supplemental table. 1: Profile of cytokines, interferons and chemokines secreted by HTR8 cells infected with the different flaviviruses.** The supernatants of each condition were collected at different times (6, 24, 48 and 72 hpi). Each value represents the mean concentrations (pg/mL) + standard deviation of three independent experiments (****p, < 0.0001; *p, < 0.046; ns, not significant).

| Condition | hpi | Cytokines (pg/mL) | | | | | Interferons (pg/mL) | |
|---|---|---|---|---|---|---|---|---|
| | | **IL-1β** | **IL-6** | **IL-10** | **IL-15** | **TNFα** | **INF-2α** | **INF-*y*** |
| ***MOCK*** | *6* | 13 ± 0.5 | 14 ± 0.2 | 0.4 ± 0.1 | 0.6 ± 0.1 | 3 ± 0.3 | 2 ± 0.3 | 5 ± 1 |
| | *24* | 13 ± 0.1 | 54 ± 2 | 0.5 ± 0.1 | 2 ± 0.1 | 6 ± 0.3 | 4 ± 2 | 6 ± 0.2 |
| | *48* | 13 ± 0.1 | 76 ± 3 | 2 ± 0.1 | 4 ± 0.1 | 7 ± 0.1 | 8 ± 2 | 7 ± 1 |
| | *72* | 15 ± 0.1 | 192 ± 1 | 2 ± 0.1 | 15 ± 4 | 9 ± 0.1 | 11 ± 1 | 15 ± 1 |
| ***ZIKV-MR77*** | *6* | 19 ± 0.5 | 15 ± 1 | 2 ± 0.7 | 1 ± 0.1 | 19 ± 0.2 | 4.4 ± 2 | 7. ± 1 |
| | *24* | 19 ± 0.5 | 82 ± 1 | 3 ± 0.2 | 3 ± 0.5 | 22 ± 0.3 | 6 ± 3 | 10 ± 3 |
| | *48* | 21 ± 2 | 275 ± 22 | 3 ± 0.1 | 6 ± 0.6 | 61 ± 0.1 | 9 ± 3 | 32 ± 6 |
| | *72* | 25 ± 1 | 4025 ± 330 | 3 ± 0.1 | 25 ± 4 | 80 ± 0.4 | 11 ± 1 | 65 ± 8 |
| ***ZIKV-MEX*** | *6* | 14 ± 1 | 11 ± 1 | 1 ± 0.2 | 0.6 ± 0.1 | 11 ± 0.5 | 4 ± 2 | 6 ± 1 |
| | *24* | 16 ± 1 | 45 ± 1 | 3 ± 0.1 | 2 ± 0.2 | 14 ± 0.1 | 6 ± 1 | 8 ± 2 |
| | *48* | 16 ± 1 | 117 ± 1 | 3 ± 0.1 | 8 ± 1 | 22 ± 0.2 | 9 ± 3 | 20 ± 7 |
| | *72* | 19 ± 3 | 1177 ± 80 | 5 ± 0.1 | 16 ± 3 | 50 ± 0.2 | 11 ± 3 | 60 ± 9 |
| ***DENV-2*** | *6* | 13 ± 0.2 | 105 ± 7 | 2 ± 0.7 | 10 ± 1 | 9 ± 0.1 | 135 + 8 | 10 + 1 |
| | *24* | 13 ± 0.2 | 165 ± 12 | 3 ± 0.1 | 13 ± 1 | 12 ± 0.1 | 161 + 22 | 14 + 2 |
| | *48* | 15 ± 0.2 | 793 ± 38 | 10 ± 0.2 | 20 ± 1 | 22 ± 0.3 | 155 + 28 | 20 + 3 |
| | *72* | 17 ± 0.2 | 3631 ± 431 | 22 ± 3 | 34 ± 3 | 41 ± 0.6 | 123 + 13 | 109 + 3 |
| ***YFV-17D*** | *6* | 13 ± 0.2 | 30 ± 1 | 1 ± 0.5 | 0.7 ± 0.2 | 11 ± 0.4 | 41 ± 25 | 8 ± 2 |
| | *24* | 14 ± 0.2 | 94 ± 4 | 2 ± 0.1 | 2 ± 1 | 14 ± 0.1 | 56 ± 13 | 11 ± 4 |
| | *48* | 15 ± 0.2 | 540 ± 32 | 4 ± 0.5 | 7 ± 1 | 24 ± 0.3 | 137 ± 13 | 31 ± 5 |
| | *72* | 16 ± 0.2 | 2268 ± 534 | 8 ± 0.7 | 21 ± 1 | 49 ± 0.1 | 179 ± 22 | 115 ± 6 |

Supplemental table 1, cont.

| Condition | hpi | Chemokines (pg/mL) | | | | | |
|---|---|---|---|---|---|---|---|
| | | **MCP1 / CCL2** | **MIP1α / CCL3** | **MCP3 /CCL7** | **IL-8 / CXCL8** | **IP-10 / CXCL10** | **VEGF** |
| ***MOCK*** | 6 | 39 ± 3 | 0.7 ± 0.2 | 1.7 ± 0.5 | 15 ± 2 | 2 ± 0.2 | 6 ± 0.2 |
| | 24 | 242 ± 17 | 1.1 ± 0.1 | 12 ± 2 | 84 ± 6 | 6 ± 0.7 | 19 ± 3 |
| | 48 | 1014 ± 102 | 1.2 ± 0.2 | 30 ± 1 | 193 ± 21 | 14 ± 3 | 29 ± 0.1 |
| | 72 | 6677 ± 104 | 1.3 ± 0.2 | 101 ± 9 | 466 ± 52 | 21 ± 3 | 81.4 ± 0.1 |
| ***ZIKV-MR77*** | 6 | 87 ± 10 | 1 ± 0.3 | 7 ± 1 | 19 ± 2 | 20 ± 2 | 6 ± 0.2 |
| | 24 | 299 ± 7 | 1.2 ± 0.4 | 18 ± 1 | 102 ± 1 | 38 ± 5 | 12 ± 2 |
| | 48 | 8185 ± 329 | 1.3 ± 0.4 | 95 ± 3 | 256 ± 36 | 73 ± 8 | 55 ± 3 |
| | 72 | 8825 ± 112 | 1.4 ± 0.5 | 352 ± 32 | 3533 ± 306 | 158 ± 12 | 234 ± 2 |
| ***ZIKV-MEX*** | 6 | 34 ± 3 | 0.8 ± 0.1 | 5 ± 0.2 | 24 ± 4 | 2 ± 0.3 | 6 ± 0.2 |
| | 24 | 255 ± 21 | 1.1 ± 0.4 | 10 ± 1 | 115 ± 16 | 6 ± 0.2 | 13 ± 2 |
| | 48 | 7561 ± 269 | 1.3 ± 0.2 | 39 ± 1 | 199 ± 19 | 12 ± 0.5 | 24 ± 4 |
| | 72 | 8244 ± 324 | 1.3 ± 0.3 | 405 ± 15 | 3036 ± 483 | 56 ± 0.5 | 60 ± 4 |
| ***DENV-2*** | 6 | 242 ± 40 | 0.9 ± 0.2 | 19 ± 0.2 | 58 ± 13 | 6 + 1 | 266 + 10 |
| | 24 | 534 ± 30 | 1.1 ± 0.3 | 28 ± 0.5 | 198 ± 42 | 12 + 1 | 272 + 3 |
| | 48 | 7684 ± 126 | 2.8 ± 0.2 | 218 ± 26 | 2815 ± 463 | 25 + 2 | 527 + 104 |
| | 72 | 8613 ± 341 | 19 ± 0.2 | 330 ± 70 | 4119 ± 631 | 36 + 4 | 1002 + 22 |
| ***YFV-17D*** | 6 | 70 ± 6 | 0.9 ± 0.1 | 2 ± 0.1 | 15 ± 4 | 7 ± 0.4 | 7 ± 0.2 |
| | 24 | 118 ± 19 | 1.3 ± 0.1 | 15 ± 0.3 | 81 ± 3 | 13 ± 1 | 11 ± 2 |
| | 48 | 7300 ± 600 | 2.4 ± 0.4 | 152 ± 7 | 504 ± 103 | 1498 ± 32 | 19 ± 3 |
| | 72 | 8395 ± 270 | 20 ± 1.9 | 523 ± 13 | 4145 ± 235 | 3650 ± 450 | 451 ± 89 |

**Supplemental table. 2: Profile of cytokines, interferons and chemokines secreted by M2-MØ infected with the different flaviviruses.** The supernatants of each condition were collected at different times (6, 24, 48 and 72 hpi). Each value represents the mean concentrations (pg/mL) + standard deviation of three independent experiments (****p < 0.0001; ***p < 0.0008; **p < 0.0056; *p, < 0.0461; ns, not significant).

| Condition | hpi | Cytokines (pg/mL) | | | | | Interferons (pg/mL) | |
|---|---|---|---|---|---|---|---|---|
| | | **IL-1β** | **IL-6** | **IL-10** | **IL-15** | **TNFα** | **INF-2α** | **INF-*y*** |
| ***MOCK*** | 6 | 49 ± 7 | 3 ± 0.2 | 21 ± 1 | 0.75 ± 0.06 | 115 ± 17 | 0.4 ± 0.1 | 0.9 ± 0.4 |
| | 24 | 103 ± 5 | 9 ± 1 | 52 ± 4 | 1.10 ± 0.11 | 389 ± 18 | 1.4 ± 0.1 | 0.9 ± 0.4 |
| | 48 | 204 ± 11 | 20 ± 1 | 89 ± 7 | 2.50 ± 0.10 | 766 ± 30 | 3 ± 0.1 | 1.1 ± 0.2 |
| | 72 | 300 ± 2 | 34 ± 2 | 123 ± 23 | 3.38 ± 0.13 | 910 ± 7 | 4 ± 0.6 | 1.2 ± 0.2 |
| ***ZIKV-MR77*** | 6 | 68 ± 3 | 12 ± 2 | 155 ± 27 | 4 ± 0.3 | 387 ± 19 | 21 ± 0.5 | 1.4 ± 0.4 |
| | 24 | 154 ± 9 | 19 ± 2 | 332 ± 74 | 6 ± 0.4 | 581 ± 36 | 21 ± 0.3 | 3.2 ± 0.2 |
| | 48 | 433 ± 74 | 35 ± 5 | 438 ± 48 | 13 ± 0.2 | 3998 ± 82 | 22 ± 0.3 | 3.4 ± 0.2 |
| | 72 | 1125 ± 137 | 71 ± 2 | 1029 ± 16 | 23 ± 0.4 | 5338 ± 282 | 22 ± 0.9 | 3.8 ± 0.3 |
| ***ZIKV-MEX*** | 6 | 24 ± 3 | 6 ± 1 | 28 ± 1 | 3.4 ± 1 | 311 ± 85 | 21 ± 0.2 | 1.3 ± 0.1 |
| | 24 | 79 ± 18 | 18 ± 2 | 183 ± 62 | 6.4 ± 1 | 676 ± 139 | 31 ± 0.7 | 1.8 ± 0.1 |
| | 48 | 222 ± 10 | 26 ± 3 | 415 ± 40 | 11 ± 0.3 | 4025 ± 984 | 31 ± 1 | 2.2 ± 0.1 |
| | 72 | 417 ± 8 | 48 ± 2 | 762 ± 94 | 20 ± 1 | 5350 ± 450 | 32 ± 2 | 2.2 ± 0.1 |
| ***DENV-2*** | 6 | 22 ± 1 | 10 ± 0.2 | 30 ± 5 | 7 ± 1 | 90 ± 7 | 57 ± 1 | 4.1 ± 0.1 |
| | 24 | 89 ± 8 | 27 ± 1 | 131 ± 5 | 12 ± 2 | 125 ± 7 | 61 ± 1 | 11 ± 0.7 |
| | 48 | 235 ± 9 | 48 ± 2 | 340 ± 10 | 17 ± 1 | 537 ± 106 | 76 ± 0.5 | 19 ± 0.3 |
| | 72 | 624 ± 32 | 72 ± 4 | 415 ± 35 | 22 ± 0.5 | 5690 ± 451 | 129 ± 3 | 25 ± 1.7 |
| ***YFV-17D*** | 6 | 186 ± 1 | 36 ± 4 | 197 ± 7 | 4 ± 0.3 | 331 ± 28 | 48 ± 8 | 3.7 ± 0.4 |
| | 24 | 248 ± 23 | 47 ± 2 | 310 ± 34 | 11 ± 1 | 657 ± 95 | 65 ± 6 | 15 ± 1.5 |
| | 48 | 404 ± 12 | 64 ± 0.2 | 357 ± 16 | 17 ± 0.3 | 1148 ± 188 | 68 ± 4 | 27 ± 1.5 |
| | 72 | 583 ± 41 | 71 ± 1 | 833 ± 85 | 21 ± 0.4 | 6445 ± 1045 | 93 ± 14 | 34 ± 2 |

Supplemental table 2, cont.

| Condition | hpi | Chemokines (pg/mL) | | | | | |
|---|---|---|---|---|---|---|---|
| | | MCP1 / CCL2 | MIP1α / CCL3 | MCP3 /CCL7 | IL-8 / CXCL8 | IP-10 / CXCL10 | VEGF |
| ***MOCK*** | 6 | 3712 + 109 | 399 + 8 | 726 + 53 | 3236 + 97 | 42 + 5 | 705 + 153 |
| | 24 | 5017.5 + 265 | 1119 + 8 | 1554.5 + 179.5 | 4068 + 38 | 52 + 4 | 812 + 72 |
| | 48 | 5876.5 + 137 | 1619 + 4 | 2877.5 + 105.5 | 4718 + 113 | 113 + 3 | 686 + 8 |
| | 72 | 7155 + 1409 | 2061 + 9 | 4566 + 81 | 5437 + 197 | 138 + 10 | 1621 + 51 |
| ***ZIKV-MR77*** | 6 | 5071 + 577 | 830 + 40 | 1400 + 100 | 3847 + 202 | 193 + 14 | 1846 + 135 |
| | 24 | 6819 + 241 | 2110 + 220 | 6822 + 764 | 4900 + 76 | 292 + 80 | 2579 + 191 |
| | 48 | 13076 + 9711 | 2305 + 75 | 8455 + 12 | 6491 + 89 | 432 + 17 | 3923 + 174 |
| | 72 | 47760 + 7464 | 3183 +88 | 10555 + 685 | 10113 + 303 | 552 + 62 | 5979 + 593 |
| ***ZIKV-MEX*** | 6 | 5581 + 601 | 843 + 53 | 5402 + 1246 | 4691 + 93 | 49 + 3 | 169 + 31 |
| | 24 | 7397 + 210 | 2129 + 384 | 6965 + 135 | 4958 + 55 | 128 + 7 | 1474 + 3 |
| | 48 | 9674 + 149 | 2754 + 74 | 10403 + 628 | 6001 + 551 | 235 + 11 | 1926 + 66 |
| | 72 | 11550 + 650 | 3273 + 81 | 12695 + 965 | 6848 + 101 | 324 + 12 | 4078 + 619 |
| ***DENV-2*** | 6 | 5753 + 368 | 427 + 5 | 1558 + 250 | 4781 + 139 | 72 + 6 | 135 + 1 |
| | 24 | 6342 + 133 | 944 + 34 | 1914 + 31 | 5667 + 501 | 105 + 1 | 171 + 26 |
| | 48 | 9012 + 99 | 1727 + 10 | 1963 + 27 | 6207 + 419 | 128 + 21 | 573 + 18 |
| | 72 | 9899 + 48 | 2403 + 65 | 3581 + 344 | 7502 + 372 | 159 + 1 | 1856 + 58 |
| ***YFV-17D*** | 6 | 4882 + 526 | 1073 + 27 | 1132 + 206 | 3255 + 95 | 186 + 50 | 1651 + 108 |
| | 24 | 6464 + 695 | 1665 + 11 | 1560 + 121 | 3737 + 47 | 298 + 14 | 1890 + 10 |
| | 48 | 7911 + 49 | 1995 + 1 | 2694 + 1077 | 4003 + 1 | 353 + 34 | 2186 + 155 |
| | 72 | 9784 + 569 | 2483 + 78 | 5827 + 1736 | 5787 + 257 | 413 + 4 | 3751 + 711 |
